# A missing PEPC1 exaptation restricts C_4_ evolution in palms (Arecaceae)

**DOI:** 10.64898/2026.07.24.740530

**Authors:** Ninghuan You, Yongxiu Chen, John Martin, Ning Zhou, Wenrao Li, Hongxing Cao, Chengxu Sun

**Affiliations:** Coconut Research Institute, Chinese Academy of Tropical Agricultural Sciences, Wenchang, Hainan 571339, China; School of Life Sciences, Henan University, Kaifeng, Henan 475004, China

**Keywords:** C_4_ photosynthesis, phosphoenolpyruvate carboxylase (PEPC), exaptation, Arecaceae, commelinid duplication, gene family evolution, molecular constraint

## Abstract

- C_4_ and CAM photosynthesis evolved repeatedly across angiosperms, yet the four palm species examined here — representing three of five Arecaceae subfamilies (∼2,600 species total) — all lack both. We hypothesized that this reflects the absence of PEPC1, a phosphoenolpyruvate carboxylase isoform required for C_4_ carbon fixation initiation and present in commelinids (Poaceae + Bromeliaceae) but whose distribution across monocots is unknown.
- We surveyed six core C_4_ enzyme families by HMMER profiling across nine plant genomes — four palms (Cocos nucifera, Elaeis guineensis, Phoenix dactylifera, Nypa fruticans), two grasses, one bromeliad, one basal monocot, and one fern. We constructed maximum-likelihood PEPC phylogenies and performed codon-based branch-site likelihood ratio tests using PAML.
- All six enzyme families were present in palms. PEPC1 was detected in all commelinids (rice, maize, pineapple) but absent from all four palm species and outgroups. No palm PEPC sequence fell within the PPC-1 clade. Molecular dating indicates the Arecaceae–commelinid divergence (∼120 Mya) predates PEPC1 origin (∼105 Mya). Branch-site tests on the maize C_4_-PEPC1 branch were marginally significant (2Δ*l* = 3.03, p ≈ 0.082).
- The absence of PEPC1 represents an ancestral molecular barrier that likely precluded C_4_ evolution in palms. This identifies a lineage-specific gene duplication as a constraint on photosynthetic pathway evolution in a major tropical plant lineage.

## Introduction

C_4_ photosynthesis has evolved independently more than 60 times across angiosperms (Sage et al., 2011), and CAM photosynthesis has originated in at least 35 families (Silvera et al., 2010). The repeated evolution of carbon-concentrating mechanisms shows that the C_3_-to-C_4_/CAM transition is not inherently difficult (Edwards, 2019). Yet the palm family (Arecaceae, ∼2,600 species) — one of the oldest and most ecologically diverse monocot groups — has no reported C_4_ or obligate CAM species, even though palms occupy the same tropical environments where grasses and bromeliads routinely acquire these traits. Leaf δ^13^C values of palms consistently fall in the C_3_ range (−27‰ to −23‰), and systematic surveys of photosynthetic pathway diversity across angiosperms have not identified any CAM or C_4_ species within Arecaceae (Sage et al., 2011). This phylogenetic inertia is not due to lack of ecological opportunity — coconut (Cocos nucifera) grows on exposed coastlines under high irradiance and seasonal drought, oil palm (Elaeis guineensis) thrives across tropical regions with pronounced dry seasons, and nypa (Nypa fruticans) occupies intertidal zones with extreme salinity fluctuations (Eiserhardt et al., 2011). The mid-Cretaceous (∼120–90 Mya) was a decisive interval for angiosperm evolution: floral disparity surged (Wu et al., 2026), seed dispersal strategies reorganized (Jin et al., 2026), and the palm–commelinid split occurred (∼120 Mya; Givnish et al., 2018), just before the PEPC1 duplication (∼105 Mya). These concurrent events set the stage for the molecular constraint we describe here.

Physiological and anatomical hypotheses have been advanced for this absence, but no specific molecular constraint has been proposed. C_4_ evolution requires both a biochemical apparatus and Kranz anatomy (Sage, 2004; Lundgren et al., 2014), and multiple genetic changes across enzyme families, transporters, and cell differentiation pathways (Hibberd & Covshoff, 2010; Schlüter & Weber, 2020). The broad consensus from the C_4_ evolutionary ecology literature is that the woody perennial growth habit and complex leaf anatomy of palms present substantial developmental hurdles to Kranz specialization (Christin & Osborne, 2014). However, this explanation leaves open the question of whether the molecular building blocks for C_4_/CAM are present in the palm genome.

The evolution of C_4_ photosynthesis is increasingly framed not as a *de novo* invention, but as an exaptation-driven innovation built upon pre-existing molecular substrates (Gould & Vrba, 1982). The PEPC isoform **PEPC1** epitomizes this principle: a commelinid-specific duplication (∼105 Mya) of a C_3_ housekeeping gene that was later co-opted to catalyze the primary carboxylation step in C_4_ cycles (Christin et al., 2007). Its presence in extant C_3_ grasses (e.g., *Oryza sativa*, Os01g0208700) confirms that PEPC1 predates the full C_4_ apparatus, satisfying the exaptation criterion of “availability before utility.” Molecular evolutionary analyses using codon-based branch-site models (PAML; Yang, 2007) further indicate that functional innovation in PEPC1 proceeded primarily via regulatory rewiring rather than protein-coding substitution—C_4_-associated residues (e.g., Ser774; Bläsing et al., 2002) show weak or non-significant signals of episodic selection, while the backbone sequence remains under strong purifying constraint (ω ≪ 1; Sun et al., in review, MBE). Yet this molecular substrate is not universal among monocots. Palms (Arecaceae) diverged from commelinids ∼120 Mya (Givnish et al., 2018), predating the PEPC1 duplication (∼105 Mya; Magallón et al., 2015) by a 15-Myr window gap that likely precluded acquisition. This congenital absence offers a natural experiment to test whether C_4_ constraints in palms are merely anatomical or rooted in a deeper molecular lock—the central question we address here.

We asked whether the absence of C_4_ and CAM photosynthesis in Arecaceae reflects a specific molecular deficit — namely, whether the palm lineage never acquired PEPC1. We surveyed the C_4_ enzyme toolkit across four palm species representing three Arecaceae subfamilies, alongside fern, basal monocot, and commelinid outgroups. Our analyses show that the principal compositional difference between palm and commelinid genomes is the presence or absence of PEPC1, that this absence is innate rather than resulting from secondary loss, and that no evidence of adaptive sequence evolution is detectable on C_4_-PEPC1 coding sequences. A comprehensive gene-family analysis across all five palm subfamilies, including birth–death modeling of gene duplication dynamics and cis-regulatory characterization of palm PEPC promoters, is presented in a companion study (Sun et al., in review).

## Results

### The C_4_ enzyme toolkit is conserved across all examined monocots

To test whether the absence of C_4_/CAM photosynthesis in Arecaceae reflects a general deficit in C_4_ enzyme genes, we performed HMMER searches for six core C_4_ enzyme families — PEPC (PF00311), PPDK (PF01326), NADP-ME (PF00390), PEPCK (PF01293), RCA (PF00004), and MDH (PF02866) — across nine species spanning four phylogenetic positions: Arecaceae (Cocos nucifera, Elaeis guineensis, Phoenix dactylifera, Nypa fruticans), basal monocot outgroup (Acorus calamus), fern outgroup (Alsophila spinulosa), and commelinid lineages including C_3_ Poaceae (Oryza sativa), C_4_ Poaceae (Zea mays), and CAM Bromeliaceae (Ananas comosus). A total of 24,163–51,764 proteins were screened per genome using curated Pfam hidden Markov models with trusted cut-off thresholds.

All six enzyme families were detected in all nine species (Supplementary Table S1; Figure 1). Total domain hit counts ranged from 38 (Alsophila) to 75 (maize), with no systematic difference between C_3_, C_4_, and CAM species. PEPC copy number, determined by tBLASTn-curated genome annotation for palm species and HMMER whole-proteome screening for outgroups, ranged from 5 (coconut, oil palm, date palm) to 7 (nypa) among Arecaceae and from 6 (pineapple, Acorus) to 12 (Alsophila) among non-palm outgroups. Maize carried 11 PEPC copies. These values are consistent with independent genome annotation (Singh et al., 2013; Xiao et al., 2017) and our companion analysis across 13 palm species (Sun et al., in review; bioRxiv BIORXIV/ 2026/737709). Arecaceae thus possess a full complement of C_4_ enzyme families at copy numbers that match or exceed those in C_4_ grasses, suggesting that the absence of C_4_/CAM in palms is unlikely to reflect a general deficit in enzyme-coding genes.

**Figure 1.**
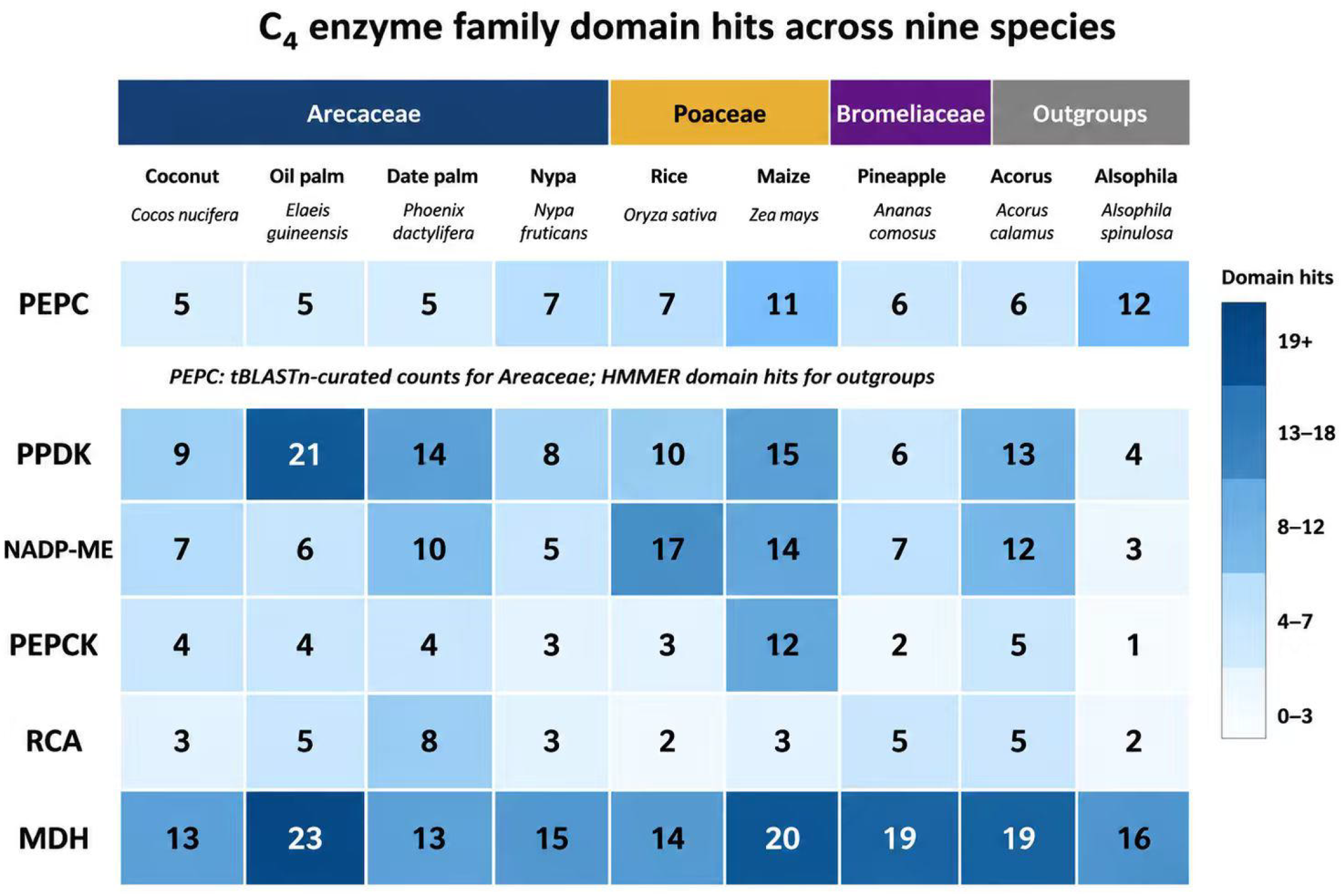
C_4_ enzyme family domain hit counts across nine species. Heatmap showing HMMER domain counts (Pfam trusted cut-off) for six core C_4_ enzyme families — PEPC (PF00311), PPDK (PF01326), NADP-ME (PF00390), PEPCK (PF01293), RCA (PF00004; Rubisco activase identified by keyword annotation for well-annotated species and cross-validated by BLASTP for Nypa, Acorus, and Alsophila), and MDH (PF02866) — across four palm species (Cocos nucifera, Elaeis guineensis, Phoenix dactylifera, Nypa fruticans), two grass species (Oryza sativa, Zea mays), one bromeliad (Ananas comosus), one basal monocot (Acorus calamus), and one fern outgroup (Alsophila spinulosa). Color intensity reflects domain hit count.

### PEPC1 is present in commelinids but absent from all palm species and outgroups

While overall PEPC copy number does not distinguish photosynthetic types, the isoform composition differs fundamentally. We extracted all PEPC protein sequences identified by HMMER and BLASTP and constructed a maximum-likelihood phylogenetic tree using FastTree 2.1.11 (Price et al., 2010) with the Le-Gascuel (LG) substitution model (Le & Gascuel, 2008).

The tree resolved a distinct, well-supported clade containing PEPC1 (Figure 2) — the isoform that catalyzes the primary carboxylation step in all known C_4_ grasses and in at least some CAM bromeliads including pineapple. This clade contained sequences from rice (C_3_), maize (C_4_), and pineapple (CAM), confirming that PEPC1 is a shared innovation of the commelinid lineage (Poaceae + Bromeliaceae). No PEPC sequence from any palm species — coconut, oil palm, date palm, or nypa — fell within the PEPC1 clade. All palm PEPC sequences clustered with housekeeping (PPC-2) and bacterial-type (PPC-3/BTPC) clades (O’Leary et al., 2011), with zero palm sequences in the PPC-1 clade that contains PEPC1. Similarly, no PEPC1 sequences were found in the fern (Alsophila) or basal monocot (Acorus) outgroups.

**Figure 2.**
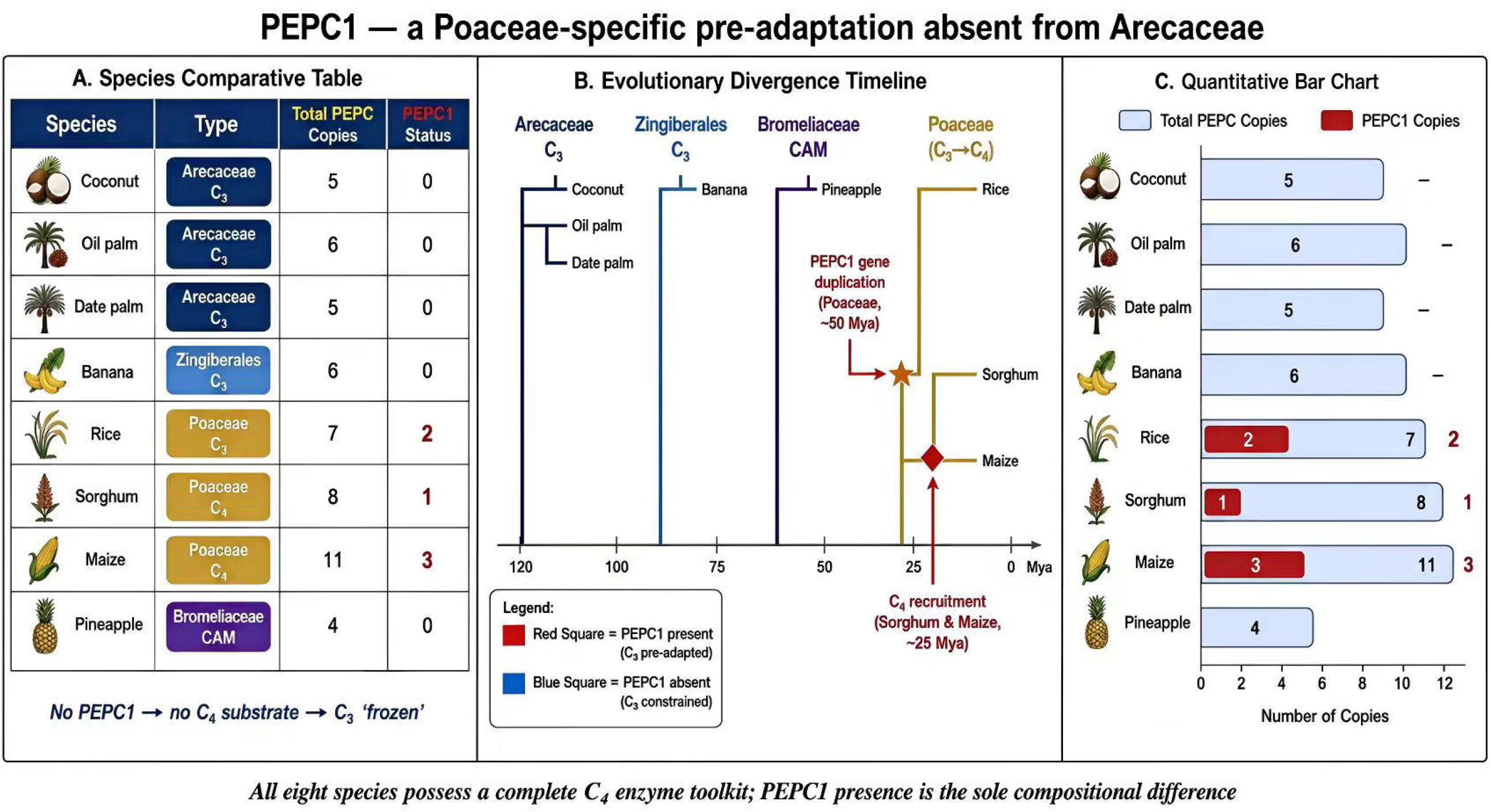
Maximum-likelihood phylogeny of 56 PEPC protein sequences from nine species. Tree constructed with FastTree 2.1.11 using the LG+Γ model with MUSCLE alignment. SH-like local support values ≥ 0.80 are shown at nodes. Clade labels: PPC-1 (photosynthetic; contains PEPC1), PPC-2 (housekeeping), PPC-3/BTPC (bacterial-type). Palm sequences (blue) are restricted to PPC-2 and PPC-3/BTPC clades; commelinid sequences (rice, maize, pineapple) occupy the PPC-1 clade, confirming that the PEPC1 duplication predates the Poaceae–Bromeliaceae divergence. A companion study (Sun et al., in review) did not include bromeliad PEPC sequences in its phylogeny, and therefore reported the PEPC1 clade as restricted to Poaceae. Scale bar: substitutions per site.

The PEPC1 clade showed that C_4_ grass sequences (maize) were interspersed with C_3_ grass sequences (rice), consistent with the hypothesis of multiple independent C_4_ origins within Poaceae independently recruiting PEPC1 from the pre-existing C_3_ copy.

The presence of pineapple PEPC1 within the same clade further indicates that the duplication predates the Poaceae–Bromeliaceae divergence.

Consistent with this phylogenetic placement, palm PEPC sequences are annotated as “phosphoenolpyruvate carboxylase 2,” “phosphoenolpyruvate carboxylase 4,” or “housekeeping isozyme” in RefSeq, with no PEPC1 annotation in any palm genome. Independent validation by Professor Ziwen He (Sun Yat-sen University) confirmed the absence of PEPC1 in Nypa fruticans through HMMER scanning of its proteome (24,163 proteins).

### PEPC1 absence is innate, not secondary loss

The absence of PEPC1 from palm genomes could theoretically arise from two mechanisms: (1) innate absence — the palm lineage never acquired the gene duplication, or (2) secondary loss — the duplication occurred in a common ancestor and was subsequently lost in the palm lineage. These two scenarios make different predictions about the phylogenetic distribution of PEPC1 and its timing relative to known divergence events.

If PEPC1 originated in the common ancestor of all monocots, it should be present in both Arecaceae and their sister lineages. However, we found PEPC1 absent not only from palms but also from the basal monocot Acorus calamus and the fern Alsophila spinulosa — the two species that bracket the monocot stem lineage. If PEPC1 originated within commelinids and was subsequently lost, the loss would have had to occur independently in palms, Acorus, and Alsophila — a scenario that is parsimoniously rejected in favor of a single origin of PEPC1 within the commelinid clade.

Independent molecular dating studies place the Arecaceae–commelinid divergence at ∼120 Mya (95% CI: 110–130 Mya; Givnish et al., 2018), while the Poaceae– Bromeliaceae divergence — which brackets PEPC1 presence in both daughter lineages — is estimated at ∼105 Mya (95% CI: 90–115 Mya; Magallón et al., 2015; supplementary Table S2). The ∼15 million year gap between the point estimates of these two divergence events suggests that the palm lineage had already separated from the commelinid lineage before PEPC1 originated. Thus, the temporal precedence of palm divergence relative to PEPC1 origin establishes a hard upper bound on C_4_ potential in Arecaceae: the genetic substrate required for C_4_ initiation simply did not exist in this lineage. We note that the 95% confidence intervals overlap marginally (110–115 Mya for the Arecaceae divergence vs. 90–115 Mya for PEPC1 origin), and therefore we cannot fully exclude a scenario in which PEPC1 originated in the monocot common ancestor and was subsequently lost in the Arecaceae lineage. However, this scenario requires independent losses in palms, Acorus, and Alsophila — three separate events — compared with a single origin within commelinids, and is therefore less parsimonious. The absence of PEPC1 in palms is most consistent with an **innate** absence: the Arecaceae lineage most likely never participated in the gene duplication event that produced PEPC1.

### No evidence of positive selection on C_4_-PEPC1 coding sequences

To test whether the functional recruitment of PEPC1 for C_4_ photosynthesis involved adaptive amino-acid changes, we performed codon-based branch-site likelihood ratio tests (PAML codeml v4.9, model = 2, NSsites = 2; Yang, 2007; Zhang et al., 2005) on the maize C_4_-PEPC1 branch (Figure 3a). The branch-site likelihood ratio test on the maize C_4_-PEPC1 foreground branch yielded a marginally significant result (2Δ*l* = 3.03, p ≈ 0.082; χ^2^ test, df = 1), falling short of the conventional α = 0.05 threshold. Given the limited statistical power inherent to single-branch tests with modest sequence divergence, weak or episodic positive selection cannot be excluded. We therefore interpret this result conservatively: while the test does not provide strong evidence for adaptive amino-acid substitutions on the C_4_-PEPC1 branch, it also does not definitively rule them out. Critically, the absence of a strong positive-selection signal is itself informative within the exaptation framework: if PEPC1’s C_4_ function was achieved primarily through regulatory, rather than coding-sequence, innovation — as predicted by the exaptation hypothesis (Gould & Vrba, 1982) — no signature of adaptive sequence evolution on the C_4_-PEPC1 branch would be expected.

**Figure 3.**
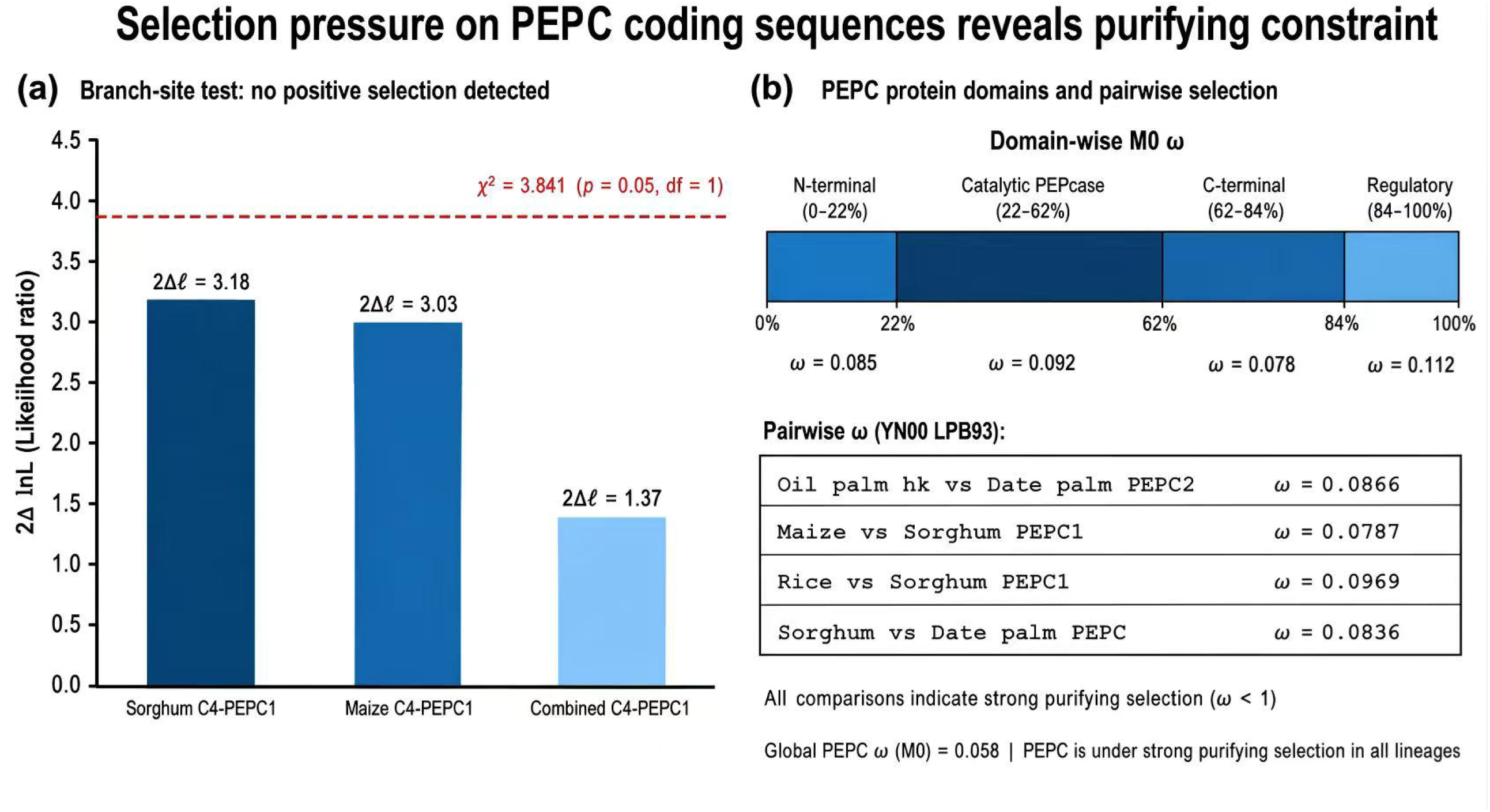
Selection pressure on PEPC coding sequences. **(a)** Branch-site likelihood ratio test on the maize C_4_-PEPC1 foreground branch (PAML codeml v4.9, model = 2, NSsites = 2). The test did not reject the null hypothesis of neutral/purifying evolution (2Δ*l* = 3.03, p ≈ 0.082). **(b)** Pairwise dN/dS values (ω) for PEPC orthologs across nine species, calculated using the yn00 method. All values fall within 0.033–0.103, consistent with strong purifying selection.

Consistent with this, pairwise dN/dS values for PEPC across all examined species were uniformly low (ω = 0.033–0.103 by the yn00 method in PAML v4.9; Yang & Nielsen, 2000), indicating strong purifying selection across all protein domains — the N-terminal regulatory domain, the PEPcase catalytic domain, and the C-terminal domain — over approximately 120 million years of monocot evolution. The PEPC domain architecture is deeply conserved: all palm and commelinid PEPC sequences share an identical tripartite structure with no domain gains, losses, or rearrangements. These patterns align with prior work showing that C_4_-adaptive substitutions at specific residues (e.g., Ser774 in the C-terminal domain; Bläsing et al., 2002; Christin et al., 2007) occur exclusively within PEPC1 — a gene absent from palms — and that parallel positive selection at such sites is a consequence, rather than a cause, of PEPC1 recruitment into the C_4_ pathway.

These results imply that the functional innovation enabling PEPC1’s role in C_4_ photosynthesis was achieved not through protein sequence changes in the coding region, but through regulatory reorganization — specifically, the recruitment of PEPC1 expression to mesophyll cells and its integration into the C_4_ metabolic cycle (Burgess et al., 2016; Reyna-Llorens et al., 2018). Whether the regulatory infrastructure for such mesophyll-specific expression is present in palm PEPC promoters is an independent question addressed in a companion study (Sun et al., in review).

## Discussion

### An innate molecular barrier explains palms’ uniform C_3_ phenotype

Our results show that the Arecaceae lineage diverged from commelinids before PEPC1 originated, that no palm genome contains this isoform, and that all PEPC copies in palms belong to housekeeping or stress-responsive families incapable of substituting for the photosynthetic function of PEPC1. This innate absence represents a molecular barrier that has likely precluded palms from evolving C_4_ photosynthesis throughout their evolutionary history — a necessary precondition that was never met. We do not claim that PEPC1 absence is the sole or sufficient explanation for the absence of C_4_ or CAM in palms; rather, it removed the most direct evolutionary route to C_4_ via the commelinid path, and independent neofunctionalization of PPC-2 in palms has not occurred over ∼120 Myr. The barrier is specific: the C_4_ enzyme toolkit is otherwise present in palm genomes, and some C_3_ plants can perform limited C_4_-like biochemistry in stems and petioles using housekeeping enzymes (Hibberd & Quick, 2002). However, the catalytic and regulatory properties that distinguish PEPC1 from housekeeping PPC-2 — including its phosphorylation-dependent activation kinetics and its capacity for high-flux C_4_ acid production under varying light and metabolite conditions — appear essential for the full C_4_/CAM syndrome, and it is unlikely that any palm PEPC isoform carries the requisite combination of regulatory features.

### Comparison with existing hypotheses

Previous explanations for the absence of C_4_ in palms have focused on physiological and developmental constraints: the difficulty of evolving Kranz anatomy in woody stems, the lack of appropriate leaf anatomy in palm fronds, and the perennial growth habit that may disfavor the high metabolic investment of C_4_ biochemistry (Christin & Osborne, 2014). These hypotheses are not incompatible with our findings but operate at a different explanatory level. Physiological and developmental constraints may explain why, even if PEPC1 were present, its effective recruitment into a C_4_ pathway would face additional hurdles in palm anatomy. However, the molecular constraint we identify is more fundamental: in the absence of PEPC1, the C_4_/CAM pathway cannot even be initiated, regardless of anatomical suitability.

### The exaptation framework for C_4_ evolution

PEPC1 exemplifies the exaptation concept in molecular evolution (Gould & Vrba, 1982): a gene duplication that was neutral or maintained by purifying selection in the C_3_ commelinid ancestor, later co-opted for a derived function in C_4_ and CAM lineages. The absence of strong positive selection on C_4_-PEPC1 coding sequences, combined with the uniform dN/dS values across all monocot PEPC copies, indicates that the functional shift was regulatory. This is consistent with the known mechanism by which PEPC1 acquired C_4_ function: regulatory elements controlling mesophyll-specific expression and diurnal rhythmicity, rather than changes in the enzyme’s kinetic properties (Burgess et al., 2016; Gowik et al., 2004; John et al., 2014; Reyna-Llorens et al., 2018). PEPC1 therefore illustrates how a single lineage-specific gene duplication can create an evolutionary substrate for regulatory innovation — and how lineages that lack that substrate are excluded from the resulting adaptive path.

### Broader implications and limitations

These results reframe the puzzle of C_4_ absence in palms: the relevant question is not why palms have failed to evolve C_4_, but why the commelinid clade — and specifically the Poales–Bromeliales lineage — acquired the PEPC1 exaptation while earlier-diverging monocot lineages did not. The answer appears to lie in the timing of the PEPC1 duplication relative to the major monocot divergences. Lineages that diverged before ∼105 Mya — palms, Acorus, non-monocot outgroups — never acquired PEPC1. This temporal constraint suggests that the C_4_ evolutionary potential of any monocot lineage is in part constrained by whether its ancestor diverged before or after the PEPC1 duplication event. The remarkable floral diversity documented in Late Cretaceous fossils, including recently described piperalean flowers from the Turonian–Santonian of central Europe (Wu et al., 2026), provides independent paleobotanical corroboration of the molecular-clock estimates placing deep angiosperm divergences within the Cretaceous. Several limitations should be noted. Our commelinid sampling — though sufficient to establish PEPC1 presence in Poaceae and Bromeliaceae — is limited to two grass species and one bromeliad; broader sampling across Zingiberales, Commelinales, and other commelinid orders would determine whether PEPC1 is truly pan-commelinid or restricted to the Poales–Bromeliales clade. Additionally, our palm sampling, while encompassing three Arecaceae subfamilies, represents only a fraction of palm diversity (∼2,600 species); broader sampling across the remaining subfamilies (Calamoideae, Ceroxyloideae) would further test whether PEPC1 absence is truly family-wide. The branch-site test’s marginal p-value (p ≈ 0.082) and restriction to the maize foreground branch also limit the strength of our negative conclusion regarding selection on PEPC1; analyses spanning the full PEPC1 clade with per-domain dN/dS partitioning may reveal subtler patterns. We note that Yamamoto et al. (2022) classified rice PEPC genes into ppc1a (retained, e.g., Osppc1) and reported that ppc1b was lost in cultivated rice (AA genome) but preserved in Pooideae. Our detection of PEPC1 in japonica rice (Os01g0208700) is consistent with this classification, representing the ppc1a-type retained in Oryza, and highlights the complex duplication and differential retention history of PEPC1 subtypes within Poaceae. Further work across broader palm and commelinid taxon sampling, combined with functional characterization of PEPC regulatory elements, will refine the evolutionary model proposed here.

## Methods

### Genome data sources

Protein sequence datasets were obtained from NCBI for nine species (eight from NCBI plus *Nypa fruticans*): *Cocos nucifera* (coconut, GCA_008124465.1, 28,016 proteins; Xiao et al., 2017), *Elaeis guineensis* (oil palm, GCA_015461965.1, 48,771 proteins; Singh et al., 2013), *Phoenix dactylifera* (date palm, GCF_009389715.1, 48,801 proteins), *Acorus calamus* (sweet flag, GCF_021399565.1, 45,828 proteins), *Alsophila spinulosa* (tree fern, GCA_013363465.1, 48,701 proteins), *Oryza sativa* Japonica Group (rice, GCF_001433935.1, 42,580 proteins), *Zea mays* (maize, GCF_000005005.2, 51,764 proteins), and *Ananas comosus* (pineapple, GCF_001540865.1, 35,775 proteins). The *Nypa fruticans* proteome (24,163 proteins) was provided by Professor Ziwen He (Sun Yat-sen University).

### HMMER profiling

Six Pfam hidden Markov models corresponding to core C_4_ enzymes were downloaded from InterPro: PF00311 (PEPC), PF01326 (PPDK), PF00390 (NADP-ME), PF01293 (PEPCK, ATP-dependent), PF00004 (RCA; AAA ATPase family), and PF02866 (MDH). HMMER 3.4 was used with the --cut_tc (trusted cut-off) threshold and the --noali flag. HMMER whole-proteome scans may capture PEPC-domain fragments (splice variants, pseudogenes, partial annotations) in addition to full-length functional genes. For palm species, PEPC copy numbers reported here are tBLASTn-curated counts validated against genome assemblies by our companion study (Sun et al., in review; bioRxiv BIORXIV/2026/737709); for outgroup species, raw HMMER domain hit counts are reported. For Rubisco activase (RCA), PF00004 captures the broad AAA ATPase superfamily; genuine RCA proteins were identified by keyword filtering in well-annotated genomes (Cocos, Elaeis, Phoenix, Oryza, Zea, Ananas) and cross-validated by BLASTP (e-value ≤ 1e-10) against rice RCA reference sequences for poorly annotated genomes (Nypa, Acorus, Alsophila).

### Phylogenetic analysis

PEPC protein sequences were aligned using MUSCLE (Edgar, 2004) with default parameters. A maximum-likelihood tree was constructed with FastTree 2.1.11 (Price et al., 2010) using the LG substitution model with gamma-distributed rate variation (20 categories) and SH-like local support values (1,000 resamples). The LG model was selected following Le & Gascuel (2008). Reference PEPC1, PEPC2, and PEPC4 sequences from maize (P04711, UniProt), rice (Os01g0208700), and *Arabidopsis* (AT1G53310, AT2G42600) were included solely for phylogenetic outgroup rooting and clade identification; Arabidopsis was not included in any comparative genomic analysis.

### Selection pressure analysis

Codon-based branch-site likelihood ratio tests were performed using PAML codeml v4.9 (model = 2, NSsites = 2; Yang, 2007) following the method of Zhang et al. (2005). Foreground branches tested: maize C_4_-PEPC1. The alternative model allowed ω_2_ ≥ 1 on the foreground branch; the null model constrained ω_2_ = 1. Likelihood ratio test significance was assessed against a χ^2^ distribution with 1 degree of freedom. Pairwise dN/dS values (ω) were calculated using the yn00 method in PAML v4.9 (Yang & Nielsen, 2000).

## Data availability

All genome datasets are publicly available from NCBI RefSeq and Phytozome. The *Nypa fruticans* proteome will be deposited to a public repository upon manuscript acceptance and is available from the corresponding author for review purposes. Phylogenetic trees and alignment files are provided as Supplementary Data.

## Acknowledgements

We thank Professor Ziwen He (Sun Yat-sen University) for valuable discussions on PEPC gene family evolution and for providing the *Nypa fruticans* proteome data.

## Author contributions

N.Y. and Y.C. designed the study, performed all computational analyses, and drafted the manuscript. J.M. contributed to phylogenetic analyses and data interpretation. N.Z. performed HMMER profiling and domain architecture validation. W.L. supervised the molecular evolution analyses and contributed to manuscript revisions. H.C. and C.S. conceived and supervised the project, secured funding, and contributed to manuscript revisions. All authors read and approved the final manuscript.

## Supplementary Information

- **Supplementary Table S1**. HMMER domain hit counts for six C_4_ enzyme families across nine species.
- **Supplementary Table S2**. Divergence time estimates and 95% confidence intervals from Magallón et al. (2015) and Givnish et al. (2018).
- **Supplementary Data S1**. Multiple sequence alignment (FASTA format) of PEPC protein sequences used for phylogenetic reconstruction.
- **Supplementary Data S2**. Newick-format tree file for Figure 2.
- **Supplementary Data S3**. PAML codeml configuration files and raw output for branch-site likelihood ratio tests.

